# Influence of asymmetric microchannels in the structure and function of engineered neuronal circuits

**DOI:** 10.1101/2024.07.09.602729

**Authors:** JC Mateus, P Melo, M Aroso, B Charlot, P Aguiar

## Abstract

Understanding the intricate structure-function relationships of neuronal circuits is crucial for unraveling how the brain sustains efficient information transfer. In specific brain regions, like the hippocampus, neurons are organized in layers and form unidirectional connectivity, which is thought to help ensure controlled signal flow and information processing. In recent years, researchers have tried emulating these structural principles by providing cultured neurons with asymmetric environmental cues, namely microfluidics’ microchannels that promote directed axonal growth. Even though a few reports have claimed achieving unidirectional connectivity of *in vitro* neuronal circuits, given the lack of functional characterization, whether this structural connectivity correlates with functional connectivity remains unknown.

We have replicated and tested the performance of asymmetric microchannel designs previously reported in the literature to be successful in the promotion of directed axonal growth, as well as other custom variations. A new variation of “Arrowhead”, termed “Rams”, was the best-performing motif with a ∼76% probability per microchannel of allowing strictly unidirectional connections at 14 days in vitro. Importantly, we assessed the functional implications of these different asymmetric microchannel designs. For this purpose, we combined custom microfluidics with microelectrode array (MEA) technology to record the electrophysiological activity of two segregated populations of hippocampal neurons (“Source” and “Target”). This functional characterization revealed that up to ∼94% of the spiking activity recorded along microchannels with the “Rams” motif propagates towards the “Target” population. Moreover, our results indicate that these engineered circuits also tended to exhibit network-level synchronizations with defined directionality.

Overall, this characterization of the structure-function relationships promoted by asymmetric microchannels has the potential to provide insights into how neuronal circuits use specific network architectures for effective computations. Moreover, the here-developed devices and approaches may be used in a wide range of applications, such as disease modeling or preclinical drug screening.

## INTRODUCTION

A key goal of neuroscience is to understand how neurons’ physical connections (structural connectivity) and how they communicate with one another (functional connectivity) work together to produce the computations underlying cognition and behavior. Theoretical studies suggest that brain networks tend to maximize the number and variety of functional motifs while using a relatively limited set of structural motifs (Bassett & Sporns, 2017). While it is virtually impossible to ascertain the prevalence of a specific motif within the entire connectome, some traits, like “feed-forward” connectivity, are common across diverse brain networks. Examples include the hippocampus’s trisynaptic circuit or the neocortex pathways activated by sensory input (e.g., visual stimuli). However, the difficulty of studying these pathways *in vivo* has hampered our ability to fully understand the dynamic interplay of structure and function.

To address this challenge, researchers have engineered *in vitro* models that take advantage of the axons’ dimensions and cue-sensing capability to impose directional connectivity between neuronal populations. Even though a wide range of patterning techniques has been employed to control axon guidance *in vitro* (reviewed in (Mateus et al., 2024; Raj et al., 2021)), the simplest and most effective at the population level is microfluidics (Habibey et al., 2022). Importantly, microfluidics can be combined with complementary techniques, such as microelectrode arrays (MEA), to dissect neuronal function in long-term experiments (Y. Liu et al., 2024). Pioneered by Taylor et al., microfluidic technology allows for the compartmentalization of different neuronal populations, interconnected via high-aspect ratio microchannels that only allow axons to grow (Taylor et al., 2005). Taylor-based microfluidics have been used extensively to obtain insights regarding axon physiology and pathology (Mateus et al., 2024; Neto et al., 2016); but also to segregate neuronal populations into bidirectionally connected compartments, modeling brain circuits, such as the cortico-striatal (Virlogeux et al., 2018), cortico-hippocampal (Brofiga et al., 2022), or the hippocampus (Vakilna et al., 2021).

While most of the literature makes use of straight microchannels, a few groups have advanced the state-of-the-art by incorporating asymmetric motifs in the microchannel design. Taking advantage of axonal outgrowth edge guidance (Renault et al., 2016), these features bias outgrowth in the researcher’s desired direction. Thus, microchannels with asymmetric motifs can be used to establish feed-forward networks in a controlled environment (Habibey et al., 2022; Mateus et al., 2024). Among these motifs, “Arrowhead” (Gladkov et al., 2017; Holloway et al., 2019) and “Tesla valve”-inspired (Na et al., 2016; Winter-Hjelm et al., 2023) designs have been reported as capable of enforcing strictly unidirectional connections. However, no detailed comparison of the performance of these designs in shaping the structure and function of *in vitro* neuronal networks has yet been made. Moreover, no study has attempted to describe the directional flow of axonal information, which has proven crucial for the characterization of the structural connectivity between hippocampal subregions (Vakilna et al., 2021).

In this study, we set out to characterize the influence of different asymmetric motifs in the structure and function of *in vitro* neuronal circuits. We test previously reported, as well as new designs, on their blocking capabilities in the axonal outgrowth of rat hippocampal neurons from embryonic origin. We demonstrate the implications of using these designs on the profiles of activity recorded along the microchannels and in the ratio of forward/backward propagating spikes. Finally, we assess the directional bias in population-to-population network burst propagation when using these asymmetric microchannels.

## RESULTS

### Asymmetric microchannel designs

We set out to design motifs that could bias axonal outgrowth via edge-guidance pathfinding along microchannels. To this end, we based our choice on the most prominent designs in the current literature and tested new variants as well.

We replicated the “Arrowhead” design, which has been reported as capable of 100% unidirectional outgrowth bias at 14 days *in vitro* (DIV) after 5 motifs (Holloway et al., 2019), by creating a design that comprised 6 repeated motifs of these arrow-shaped structures (here termed “Arrows”). We also used this design as the basis to create a variant that, instead of stalling axonal outgrowth at the arrowheads’ corners, leads axons to a diverting loop that traps and re-routes axonal outgrowth. We named this design “Rams”, inspired by the shape of rams’ horns. We were also motivated by the Tesla valve concept, originally proposed by Nikola Tesla to allow fluidic control without any moving mechanical parts (Nikola Tesla, 1920), to design 2 variants that could re-route axonal outgrowth and create a unidirectional bias (“Tesla” and “Tesla v2”). These 2 variants differ in the number of motifs (8 in “Tesla”; 6 in “Tesla v2”) and the aggressiveness of the re-routing loop, being “Tesla v2” the variant with the sharpest angle and the one most similar to the original design. A recent study has since proposed a design that is very similar to “Tesla”, along with another Tesla valve-inspired design (Winter-Hjelm et al., 2023). Each design was implemented in microfluidic devices that comprised 2 compartments, interconnected by 16 microchannels for compatibility with standard 252-recording microelectrode arrays (MEAs). Scanning electron microscopy (SEM) images of the fabricated designs’ main variants can be seen in **Fig. 1A**.

**Figure 1.**
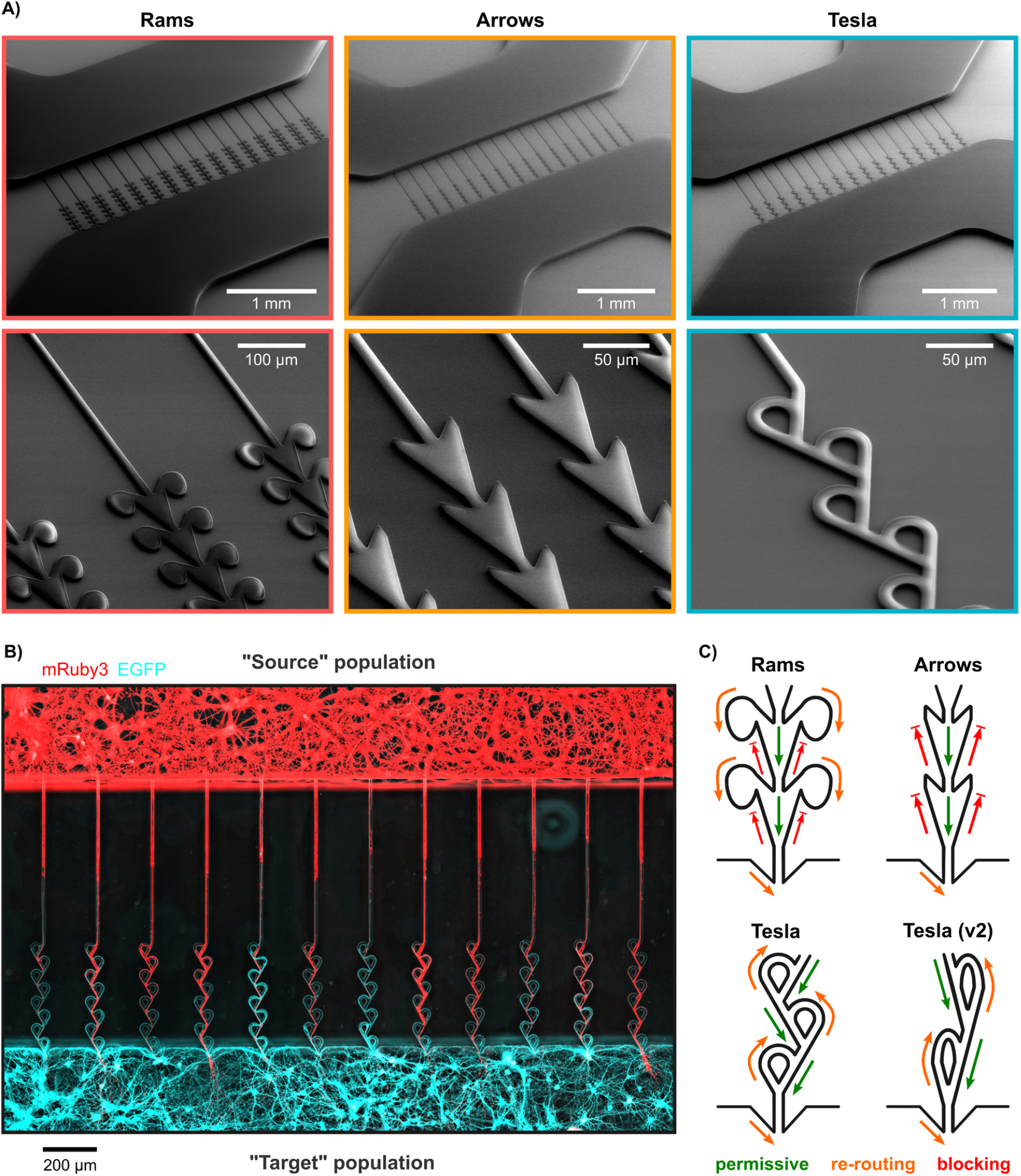
**(A)** Scanning electron microscopy (SEM) images of the fabricated designs’ main variants (“Rams; “Arrows”; “Tesla”). Low magnification images (top) encompass the entire microfluidic device design, while high magnification images (bottom) detail the respective motif design. **(B)** Widefield microscopy mosaic image (20× objective) of different neuronal populations (“Source” and “Target”) at 14 days in vitro, selectively expressing mRuby3 (red) and EGFP (cyan), cultured on a microfluidic device with the “Tesla” motif. **(C)** Schematics of the rationale for each motif design. Green, orange and red arrows indicate axonal outgrowth-permissive, re-routing and blocking capabilities, respectively.

The 4 motifs (“Rams”; “Arrows”; “Tesla”; “Tesla v2”) were employed in 1030 μm-long microchannels (measured in a straight line) that connected separate neuronal populations (“Source” and “Target”) **(Fig. 1B)**. Each microchannel comprised a straight segment (550 μm-long) to facilitate axonal outgrowth in the desired direction and an asymmetric segment (480 μm-long) to block and/or re-route axonal outgrowth in the non-desired direction. All designs kept the same microchannel height (5 μm) throughout their length. At one end of the microchannel, the compartment wall (“Target” compartment) had indentations to prevent axon growth perpendicular to the microchannels’ exit **(Fig. 1C)**, as commonly occurs in Taylor-based microfluidics (Peyrin et al., 2011). The rationale for each design is schematized in **Fig. 1C**.

### Performance of different designs in creating axonal outgrowth bias

To assess the efficacy of each design at promoting unidirectional axonal growth, we started by culturing neurons in glass coverslips and virally transducing the “Source” and “Target” populations with adeno-associated viruses (AAVs) encoding for mRuby3 and EGFP, respectively. Both viral constructs yielded a high transduction efficacy, which allowed ubiquitous expression of the different fluorescent proteins in the two neuronal populations **(Fig. 1B)**. However, despite maintaining a hydrostatic pressure gradient to hinder viral leakage between the two populations, we observed some cross-contamination of viruses that hindered formal analysis of axon blocking. Thus, to quantitatively compare the axonal outgrowth blocking probability of each microchannel design, we opted to use monocultures (“Target”-only) expressing EGFP **(Fig. 2A-B)**.

**Figure 2.**
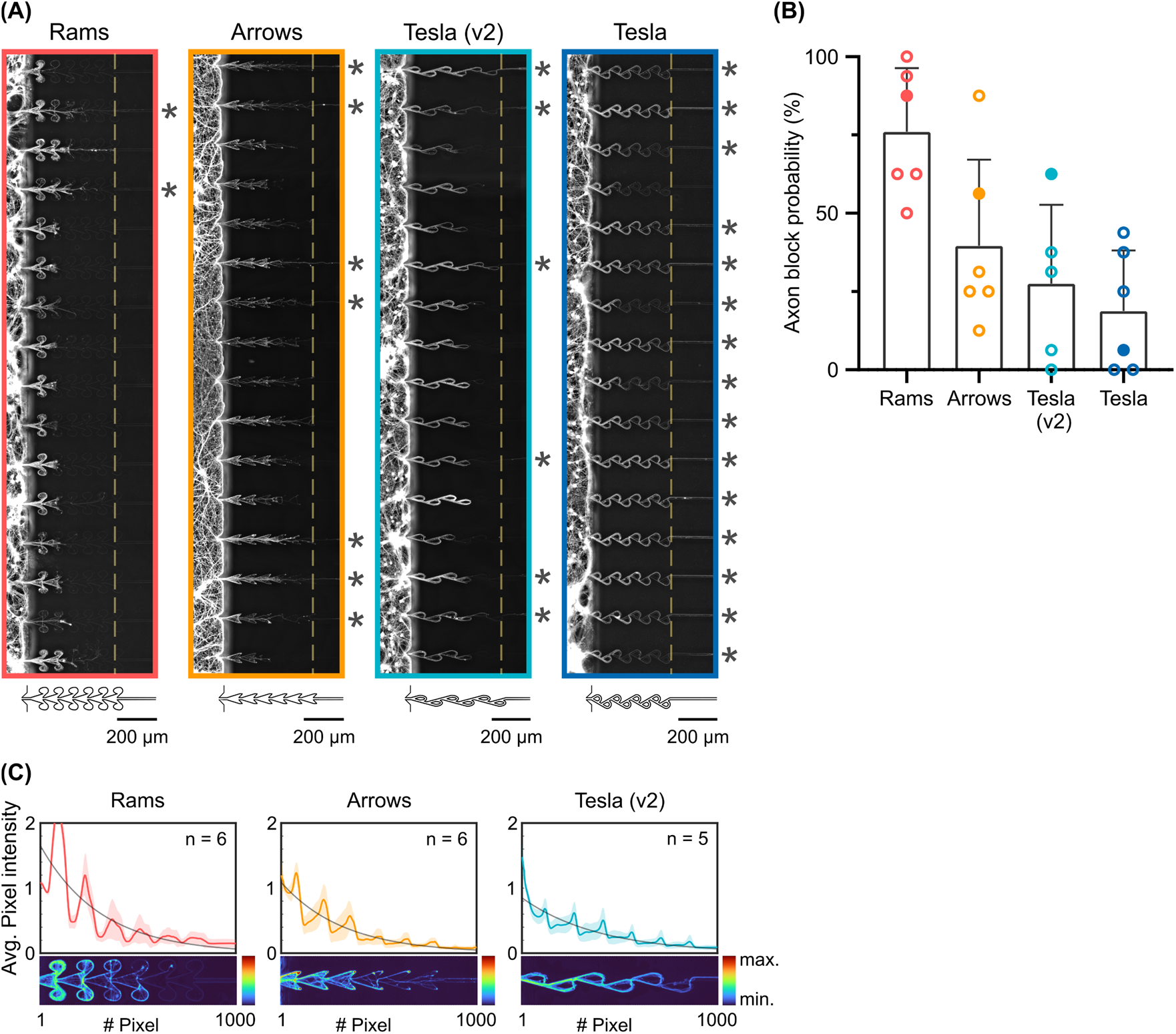
**(A)** Widefield microscopy mosaic images (20× objective) of representative neuronal monocultures (seeded on the “Target” compartment) at 14 days *in vitro* (DIV) for each design. All cultures are expressing EGFP (grayscale). Asterisks mark microchannels that failed to completely block axonal outgrowth before the straight segment of the microchannel (dashed line). **B)** Percentage of microchannels that completely block axonal outgrowth per design. Filled dots represent the cultures shown on A). **C)** Normalized average pixel intensity values along the microchannels. The average intensity profiles (and colorized shade for the SD) of 5-6 independent experiments (16 microchannels per experiment) are plotted together with a fitted exponential curve. Per design, a representative microchannel is colorized with the Turbo lookup table (LUT).

We quantified the percentage of microchannels capable of completely blocking axonal outgrowth before reaching the straight segment of the microchannel. At 14 DIV, no motif was capable of completely blocking axonal outgrowth consistently. Per order of effectiveness, the mean ±SD percentage of success in complete blocking was 76% ±20% for “Rams” (n = 6); 40% ±20% for “Arrows” (n = 6); 28% ±25% for “Tesla v2” (n = 5); and 19% ±19% for “Tesla” (n = 6) **(Fig. 2B)**.

It is important to note that while asymmetric microchannels might not completely prevent axonal outgrowth towards the “Source” population; by diverting many axons, they decrease the overall number of axons that do so. At this stage, given the small size of axons and their complex growth dynamics (e.g., ramifications), it is challenging to grasp the exact number (or fraction) of diverted axons. Still, to get a better approximation of the filtering capacity of each motif, we analyzed the fluorescence pixel intensity profiles along the better-performing designs’ microchannels (“Rams”, “Arrows” and “Tesla v2”). This analysis demonstrated that the pixel intensity decays the most along the microchannels with the “Rams” motif **(Fig. 2C)**, which provides an indirect measure of better performance at blocking axonal outgrowth with this motif.

### Asymmetry in axonal activity along the microchannel

After structural characterization of the motifs’ performance in biasing axonal outgrowth direction, we set out to characterize their influence on the function of neuronal circuits. To this end, we combined microElectrode arrays with microFluidics (µEF devices) to record neuronal activity in neuronal cultures at 12-21 DIV. The alignment compatibility with 252-recording electrode MEAs allowed for the probing of the 16 microchannels, encompassing 5 electrodes each, per experiment **(Fig. 3A)**. Importantly, since the number of microchannels matches the number of electrode columns, every microchannel is probed electrophysiologically. Thus, theoretically, every event propagating between populations (“Source” and “Target”) can be recorded. This configuration enabled the straightforward separation of activity from the “Source” and “Target” populations, as well as from the axons interconnecting them via the microchannels **(Fig. 3B)**.

**Figure 3.**
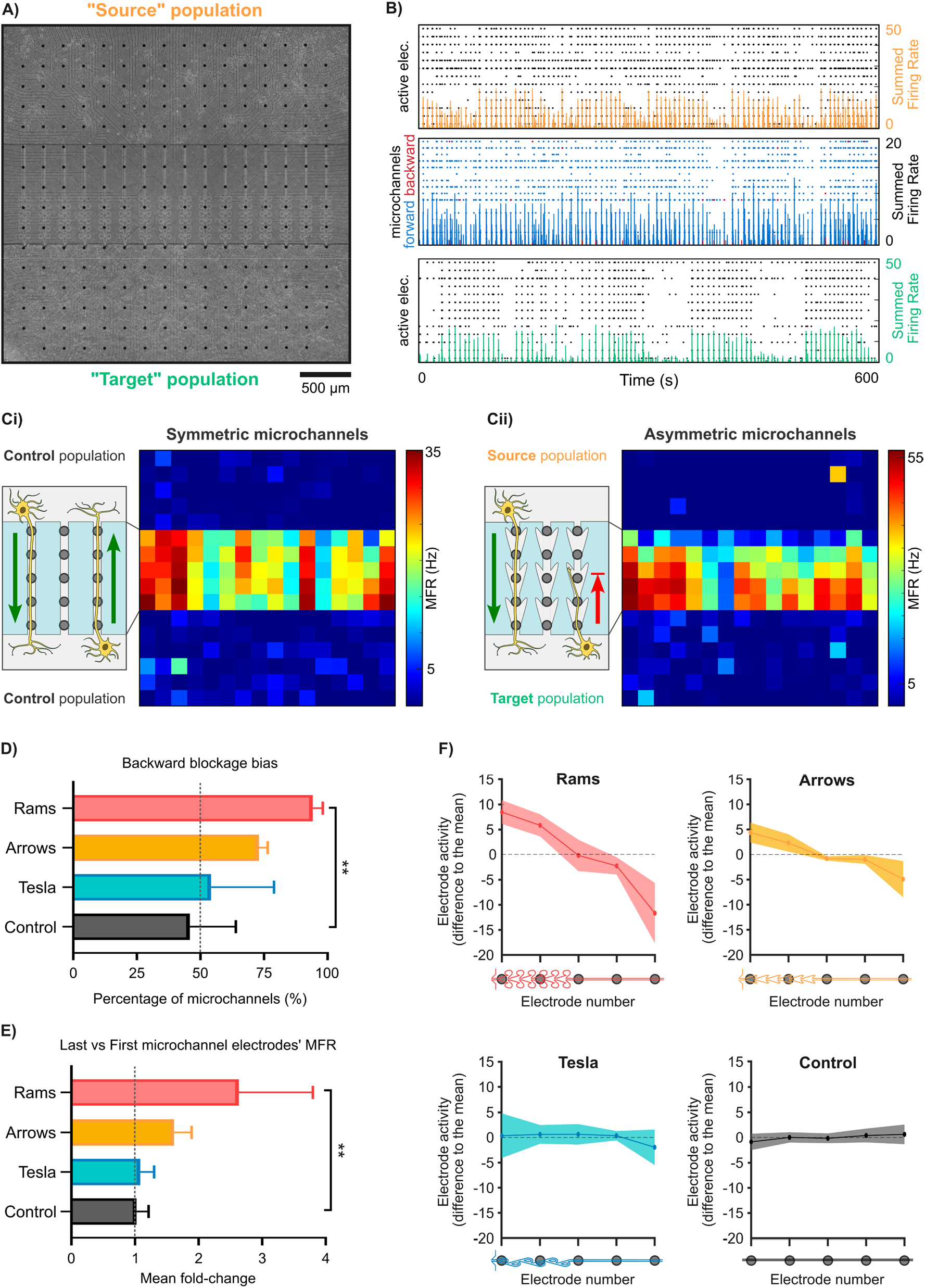
**(A)** Phase-contrast microscopy image (20× objective, mosaic) of a hippocampal co-culture at 21 days *in vitro* (DIV). The asymmetric microchannels comprise the “Rams” motif. The combination of microElectrode arrays and microFluidics (µEFs) allows the compartmentalization and probing of two neuronal populations (“Target” and “Source”), as well as the assessment of all event propagation between them. **(B)** Representative raster plot of 10 min of spontaneous activity for each of the three sections of a µEF: “Source” population; microchannels; “Target” population. The summed firing rate (color-coded) is overlayed in each plot. On the microchannels’ plot, each line represents a microchannel (i.e., one electrode per microchannel) and only events detected as propagating throughout the whole microchannel are depicted (color-coded depending on the direction as “forward” or “backward”). Culture at 12 DIV on a µEF device with the “Rams” motif. **C)** Schematics of the predicted effect of microchannels on axonal outgrowth and representative activity maps (each pixel represents an electrode’s mean firing rate (MFR)) for µEFs with **i)** symmetric microchannels (control) and **ii)** asymmetric microchannels. **D)** Horizontal bar graph showing the percentage of microchannels with a higher MFR in the asymmetric segment, when compared to the straight segment of the microchannel (i.e., with backward blockage bias), per design (n = 3-6 independent experiments, Kruskal-Wallis test with Dunn’s multiple comparisons, **p < 0.01, mean ±SD). **E)** Horizontal bar graph showing the fold-change in mean firing rate (MFR) of the two last vs two first electrodes (in the “Source”-to-”Target” direction) within the microchannels for each design (n = 3-6 independent experiments, Kruskal-Wallis test with Dunn’s multiple comparisons, **p < 0.01, mean ±SD). **F)** Average profile of activity (difference to the mean) along the 5 electrodes within the microchannels, per design (n = 3-6 independent experiments, colorized shade for the SD across recordings).

Taking advantage of this capability to record propagating axonal signals (as in (Mateus et al., 2021)), we characterized the motifs’ influence in the microchannels’ firing rate profile **(Fig. 3)** and the direction of the events’ (i.e., action potentials (AP)) propagation **(Fig. 4)**. For these and the remainder of the Results, we pooled the data from “Tesla” and “Tesla v2” experiments since we did not observe any significant differences between these designs.

**Figure 4.**
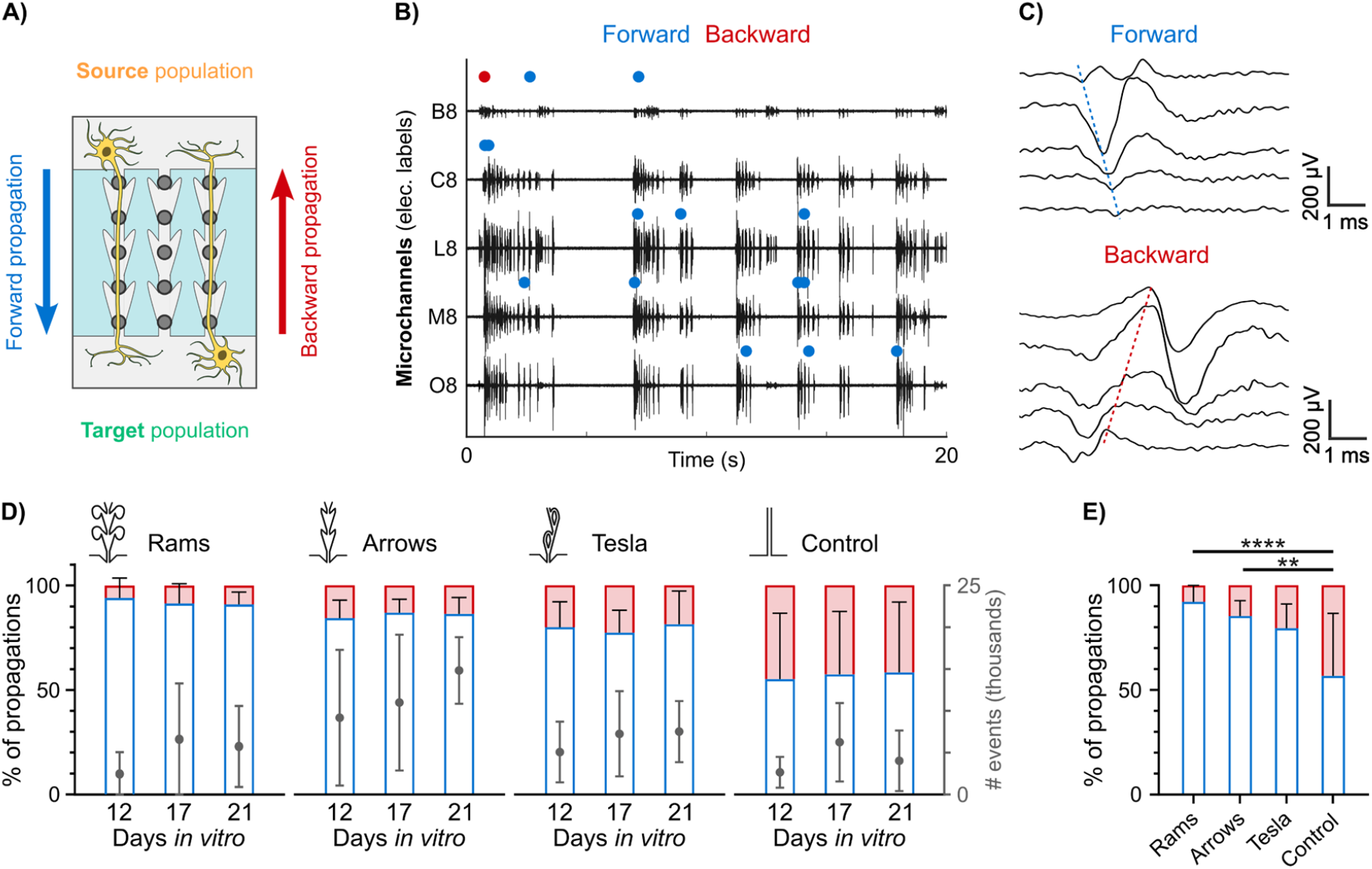
**(A)** Schematic representation of the two possible directions for signal propagation (“forward” and “backward”). **(B)** Example traces of 20 s of activity within 5 microchannels (one electrode per microchannel). Events detected as forward or backward propagations are marked by blue and red dots, respectively. **C)** Examples of forward and backward propagating events. Events were required to propagate along the whole microchannel (5 electrodes) to pass the detection criteria. **D)** Stacked bar plots showing a longitudinal analysis (12-21 days *in vitro*) of the percentage of forward (blue) and backward (red) events and the total number of detected propagating events (right y-axis), per design (n = 3-9 independent experiments, mean ±SD). **E)** Stacked bar plots comparing the percentage of forward and backward events across all recordings per design (n = 12-20, Kruskal-Wallis test with Dunn’s multiple comparisons, **p < 0.01, ****p < 0.0001, mean ±SD).

We hypothesized that, given their blocking capabilities, asymmetric microchannels should also exhibit an asymmetry in firing rate **(Fig. 3C)**. To this end, we compared the firing rate of the electrodes within the straight and asymmetric segments of the microchannels. For the “Rams” design, 94% ±4% (n = 5) of the microchannels presented higher mean firing rate (MFR) in the asymmetric segment than in the straight segment **(Fig. 3D)**. This design also presented the highest average fold-change in MFR (2.6 ±1.2) between segments **(Fig. 3E)**. This increase in MFR occurs most probably due to an accumulation of signal sources (i.e., axons) in the vicinity of the electrodes within the asymmetric segment of the microchannel, which demonstrates a backward blockage bias when using these motifs. This is also supported by the activity profiles along the 5 electrodes within the microchannels, which demonstrated a trend of increasing activity in the “Source”-to-”Target” direction **(Fig. 3F)**.

Direct measures of axonal conduction should provide a clear indication of the preferred directionality of axonal outgrowth and information flow; thus, we analyzed the directionality of AP propagation along the microchannels. We labeled as “forward” propagating events APs propagating in the “Source”-to-”Target” direction, whereas events propagating from the “Target”-to-”Source” were labeled as “backward” **(Fig. 4A-C)**.

We assessed the direction of axonal conduction during network development. These longitudinal experiments (12-21 DIV) revealed that asymmetric microchannels consistently exhibited a much higher prevalence of forward propagating events (at least 4 times more), regardless of the DIV or motif **(Fig. 4D)**. Still, the “Rams” motif exhibited the highest fraction of forward propagating events, particularly at 12 DIV (94% ±9% (n =4)). At 21 DIV, this fraction was 91% ±6% (n =4) which supports the rationale that the forward spiking bias is maintained throughout network development. Given the lack of variability in these ratios during development, we pooled the data from all DIV to compare with controls (i.e., straight microchannels). This analysis returned the “Rams” and “Arrows” motifs as performing significantly differently than controls, but not “Tesla” **(Fig. 4E)**.

It is important to note that, throughout this section’s results, we observed the same performance trend (“Rams” > “Arrows” > “Tesla”) as in the structural studies.

### Asymmetry in network burst propagation

Following the characterization of spiking activity propagation along the microchannels, we analyzed whether this activity led to a recruitment of network-level activation. Network bursts (NB) are hallmarks of neuronal synchronization *in vitro* and are often used as a proxy for assessing inter-population communication (Barral et al., 2019; DeMarse et al., 2016; Pigareva et al., 2024; Winter-Hjelm et al., 2023). Here, we used a strict causality window [20, 240] ms to identify instances of population-to-population spontaneous NB propagation. If the onset of both populations’ NB fell into this causality window, we considered this as a putative NB propagation **(Fig. 5A)**. Once again, we distinguished these propagations as “forward” and “backward” depending on the direction of propagation “Source”-to-”Target” and “Target”-to-”Source”, respectively. Importantly, we generally did not observe significant differences between the overall number of NBs detected for each population category **(Fig. 5B)**, which could bias the results.

**Figure 5.**
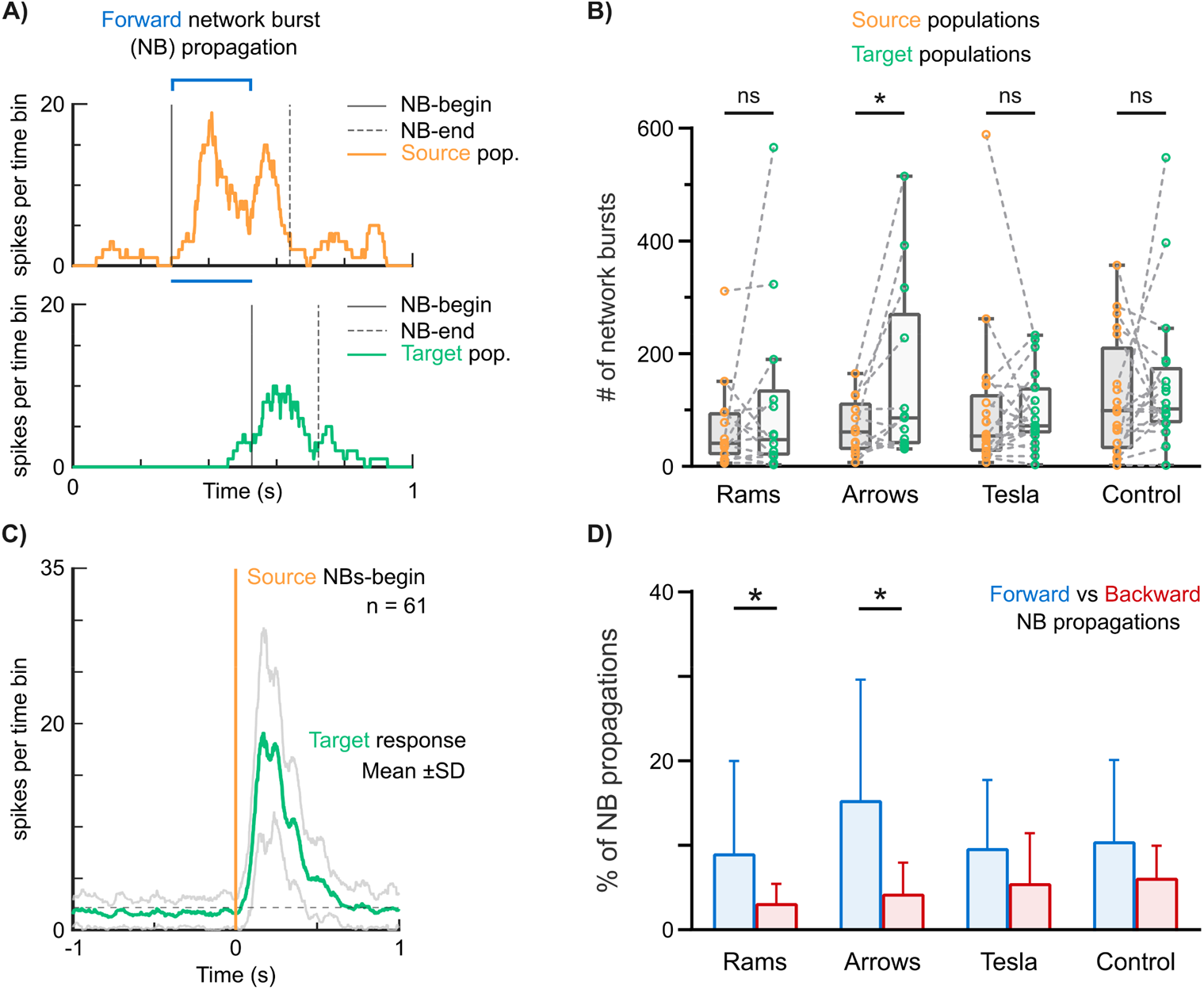
**(A)** Example of a single putative network burst (NB) propagation from a “Source” to a “Target” population (“forward” direction). The activity was recorded at 12 days *in vitro* (DIV) from populations interconnected via asymmetric microchannels (“Rams”). Traces show the summed spike train data (1 ms time bins) of each population. **(B)** Tukey plots of the total number of detected NBs per population and per design (n = 12-20, Kruskal-Wallis test with Dunn’s multiple comparisons, *p < 0.05). **C)** Event-trigger averaging of the spontaneous activity from a “Target” population after NB onset on the “Source” population. The dashed line represents the baseline activity of the “Target” population. The activity was recorded at 12 DIV from populations interconnected via asymmetric microchannels (“Arrows”). **(D)** Bar plots comparing the fraction of NB events that propagate “Source”-to-”Target” (forward direction) and “Target”-to-”Source” (backward direction), per design (n = 7-12, Kruskal-Wallis test with Dunn’s multiple comparisons, *p < 0.05, mean ±SD).

We could observe that Source’s NBs were capable of driving Target’s responses at 12 DIV on populations interconnected via asymmetric microchannels **(Fig. 5C)**. Thus, we next estimated the fraction of NBs generated in each population that propagated to its counterpart. Even though most NBs did not propagate (or elicit a proper NB), we observed that the “Rams” and “Arrows” motifs biased NB propagation to preferentially occur in the forward direction **(Fig. 5D)**. On average, the ratio between forward and backward-propagating NBs was of 2.9 and 3.6 for these motifs, respectively. These results demonstrate, once again, that these designs effectively create a directional bias in the neuronal circuits’ function. The relatively low number of microchannels may explain why most cultures maintained a high degree of population activity independence in our experiments (DeMarse et al., 2016). In the future, this number may be tuned to produce circuits with more or less structural-functional bias, as intended by the researcher.

## DISCUSSION

Here, we set out to characterize the influence of asymmetric microchannel designs in the structure and function of *in vitro* neuronal circuits. Following the principles of edge-guidance, such designs exert blocking, delaying and re-routing effects on axonal outgrowth, which we used to bias connectivity between “Source” and “Target” neuronal populations.

Even though Tesla valve-inspired designs have gained some traction in the microfluidics field (reviewed in (Purwidyantri & Prabowo, 2023)), they had not been tested in this context until very recently (Winter-Hjelm et al., 2023). Despite qualitative indications from the literature, our Tesla valve-inspired designs performed relatively poorly in the re-routing of axonal outgrowth and the establishment of directional functional connectivity. Similarly, we could not replicate previously reported results with “Arrowhead” (Holloway et al., 2019), despite our efforts (e.g., more motifs) to increase the asymmetry bias in the “Arrows” design. Ultimately, experimental details, such as cell density or viability, may underlie the observed differences. It is also worth noting that “Source” axons can serve as guiding lines to axons growing in the opposite direction (Renault et al., 2016). Since our structural experiments were performed with mono-cultures (“Target”-only), as most studies in the field, the results may be already overestimating the motifs’ structural efficiency.

“Rams”, a new motif that combines both blocking capabilities from “Arrows”-like designs and re-routing capabilities of “Tesla” or “Arches” designs (Renault et al., 2016), proved to bias structural and functional connectivity across all experiments the most. Impressively, ∼76% of the microchannels with this motif completely blocked “Target” axons at 14 DIV, and >90% of the propagating events were in the forward direction until, at least, 21 DIV. This bias in information flow was also observed in the relative fraction of NB events that propagate from the “Source” to the “Target” population, a clear indication of directional functional connectivity. Overall, our results pose the “Rams” motif as a strong candidate to be used in studies aiming to achieve a high degree of structural and functional directionality.

State-of-the-art node-based designs have demonstrated that strict unidirectional connectivity can be achieved *in vitro*. For example, a stomach-like design was reported to achieve unidirectional structural connectivity between nodes with ∼92% probability (Forró et al., 2018). Nanochannels of ∼150 nm have been shown to completely block axon growth while allowing dendritic spines to pass through, thus, in combination with microchannels that guide axon growth, can be used to establish pure feed-forward circuits (Mateus et al., 2022). These studies’ concepts can be used to form small-scale neuronal circuits (i.e., with dozens of neurons), but, intrinsically, cannot be scaled for implementation in standard Taylor-based microfluidics with hundreds of thousands of neurons. However, motif designs, such as the ones here proposed, can be easily implemented in Taylor-based microfluidics or micropatterning configurations (Yamamoto et al., 2023). If needed, their performance may be further enhanced by incorporating chemical cues (e.g., gradients of trophic factors) that attract axonal outgrowth and are commonly used in standard microfluidic devices (Rifes et al., 2020).

Finally, it is important to consider that strict unidirectional structural connectivity does not necessarily translate to function. We and others have previously shown that signal flow along axons can be spontaneously bidirectional (C. Liu et al., 2022; Mateus et al., 2021; Rózsa et al., 2023). While more research is needed to dissect the structure-function relationships that emerge in these *in vitro* models, such asymmetric microchannels may be readily used in several research fields. For example, mechanistic studies of physiology or disease, where certainty regarding axon directionality is crucial for the interpretation of the results (e.g., studies of “prion-like” mechanisms; axonal transport), could benefit greatly from the implementation of these devices (Courte et al., 2023).

## DATA AVAILABILITY STATEMENT

The data produced for this study is available from the corresponding author upon reasonable request.

## ACKNOWLEDGMENTS

This work was partially financed by FEDER - Fundo Europeu de Desenvolvimento Regional funds through the COMPETE 2020 - Operacional Programme for Competitiveness and Internationalisation (POCI), Portugal 2020, and by Portuguese funds through FCT - Fundação para a Ciência e a Tecnologia/Ministério da Ciência, Tecnologia e Ensino Superior in the framework of the projects PTDC/EMD-EMD/31540/2017 (POCI-01-0145-FEDER-031540) and PTDC/MED-NEU/28623/2017 (NORTE-01-0145-FEDER-028623). MA was supported by FCT through the Scientific Employment Stimulus (CEECIND/03415/2017).

## METHODS

### Microfluidic design and fabrication

The polydimethylsiloxane (PDMS) microfluidic chambers used in this study were designed to bias axonal outgrowth in a unidirectional manner. The “Arrows” motif mimics the design proposed by (Holloway et al., 2019), whereas the “Tesla” motif drew inspiration from the original Tesla valves proposed by Nikola Tesla in the patent #US001329559 (Nikola Tesla, 1920). The “Rams” motif is original but emerged from the “Arrows” motif.

In all microfluidic devices, the number of microchannels matched exactly the number of microelectrode columns of 256-electrode planar MEAs (16 microchannels in total). This allowed the detection of all activity propagation between the neuronal populations. Concisely, the microfluidic chambers were composed of two separate somal compartments interconnected by 16 microchannels with 1030 μm length (measured in a straight line) × 5 μm height × 10 μm width dimensions and interspaced by 200 μm. Control devices with straight microchannels with the same dimensions were used in the electrophysiology experiments.

From the microfabricated SU-8 mold, the PDMS microfluidic chambers were produced by replica molding. PDMS was prepared using a 10:1 mix of silicone elastomer and its curing agent (Sylgard 184, Dow Corning) and degassed using a vacuum desiccator. Polymerization was achieved at 70 °C for 3 h, after which the PDMS microfluidic chambers were unmolded and cut. The medium reservoirs were manually punched with a steel biopsy punch (Ø 6 mm).

### Substrate preparation

Coverslips or planar MEAs (Multi Channel Systems (MCS), Germany) of 252 titanium nitride recording microelectrodes (30 μm in diameter) and 4 internal reference electrodes (organized in a 16 by 16 square grid) were used as cell culture substrates for all experiments.

Cell culture substrates were prepared as previously detailed (Lopes et al., 2018). Briefly, both the substrate (i.e., coverslip or MEA) and the microfluidic chambers were sprayed with 70% ethanol, allowed to air-dry inside a laminar flow hood, and sterilized by ultraviolet (UV) light exposure. In the case of MEA substrates, μEF assembly was guided by a stereomicroscope, to correctly align the microchannels with the microelectrode grid. Each microchannel encompassed 5 microelectrodes, 3 within the straight segment and 2 within the asymmetric segment. Moreover, this configuration allowed for each neuronal population to be probed with a minimum of 78 microelectrodes. An image of an assembled µEF is shown in **Fig. 3A**. After alignment, the assembled devices were air plasma-cleaned for 1 min at 15W and coated with poly-D-lysine (PDL) (20 µg/ml) to promote cell adhesion. Excess coating was washed out thrice before cell seeding.

### Cell culture

Experimental procedures involving animals were carried out following current Portuguese laws on Animal Care (DL 113/2013) and the European Union Directive (2010/63/EU) on the protection of animals used for experimental and other scientific purposes. The experimental protocol (reference 0421/000/000/2017) was approved by the ethics committee of the Portuguese official authority on animal welfare and experimentation (Direção-Geral de Alimentação e Veterinária). All efforts were made to minimize the number of animals and their suffering.

Primary embryonic rat hippocampal neurons were isolated from Wistar rat embryos (E18), as in (Lopes et al., 2018). Briefly, hippocampus tissues were dissected in Hank’s Balanced Salt Solution (HBSS) and enzymatically digested in 0.6% (w/v) trypsin (1:250) in HBSS for 15 min at 37 °C. Then, trypsin was inactivated with HBSS containing 10% fetal bovine serum and washed away. Tissue fragments were mechanically dissociated with a plastic pipette and the cells’ suspension was filtered with a 40 µm strainer (Falcon) to exclude remaining tissue clumps. After cell counting, 100k viable cells suspended in 4 μl were seeded in each somal compartment of previously prepared µEFs. Cells were cultured in Neurobasal Plus medium supplemented with 0.5 mM glutamine, 2% (v/v) B27 Plus, and 1% (v/v) penicillin/streptomycin (P/S) for 21 days. Cultures were kept in a humidified incubator at 37 °C supplied with 5% CO_2_.

Transductions with adeno-associated viruses (AAVs) were performed to live-image neuronal morphology and axonal outgrowth. For neuronal morphology, scAAV-9/2-hSyn1-chI-loxP-EGFP-loxP-SV40p(A) (8.5 x 1012 vg/ml titer) or scAAV-DJ/2-hSyn1-chI-loxP-mRuby3-loxPSV40p(A) (7.2 x 10^12^ vg/ml titer) were added to each somal compartment (0.3 µl per somal compartment at 1-3 DIV). Viral vectors were produced by the Viral Vector Facility of the Neuroscience Center Zurich (Zentrum für Neurowissenschaften Zürich, ZNZ, Switzerland).

### Imaging analysis

Fluorescence imaging experiments were performed with hippocampal neurons at 14 DIV. Images were acquired by a sCMOS camera Prime 95B, 22mm (Teledyne Photometrics, UK), mounted on a Nikon Eclipse Ti2-E (Nikon, Japan) inverted microscope with a Nikon Pl Apo 20×/0.75NA objective. Image acquisition was performed using Micromanager (Version 1.4) and images were processed using ImageJ (Fiji).

Tiled images, such as the ones presented in **Fig. 2A**, were produced via stitching of the original images. Importantly, due to their size and higher levels of protein expression, somata are brighter and thus more visible than axons on microscopy images. To enhance the signal intensity of the axons compared to the somata, the Enhance Local Contrast (CLAHE) algorithm was applied to the tiled images with default parameters. For the analysis presented in **Fig. 2B**, images were further saturated to emphasize axons with low signal. Microchannels were manually labeled according to the complete blockage of axonal outgrowth by the motifs.

For the analysis presented in **Fig. 2C**, regions of interest (ROI) of defined size (encompassing all microchannels) were outlined. A background subtraction (100-pixel radius) was applied to the resulting 8-bit tiles. Then, the analysis proceeded in Matlab using custom scripts. Pixel intensity profiles were obtained for each image via column-wise averaging. Then, pixel intensity was normalized against the first 5-pixel columns (corresponding to the entrance of the microchannels), for each image. The profiles of 5-6 independent experiments (i.e., images) per design were averaged. Finally, a 20-pixel moving average was applied to the traces before plotting and fitting the exponential curve.

### Electrophysiological recordings

Hippocampal neurons at 12-21 DIV were used in electrophysiological experiments. Recording sessions started after 10 minutes of adjustment to recording conditions.

Electrophysiological recordings of spontaneous electrical activity were obtained at a sampling rate of 20 kHz for 10 minutes. All recordings were obtained using a commercial MEA2100-256 system (Multichannel Systems MCS, Germany) mounted on an incubated (37 °C) inverted widefield microscope (Axiovert 200M, Zeiss or Eclipse Ti2-E, Nikon) stage supplied with humidified 5% CO_2_. Temperature was maintained at 37 °C by external temperature controllers.

### Action potential detection and propagation characterization

Raw signals were high-pass filtered (200 Hz) and APs were detected by a threshold method set to 5× the standard deviation (SD) of the peak-to-peak electrode noise. An AP time was extracted at this surpassing point and no detection was considered for the next 3 ms (“dead time”). Analysis was then performed offline using custom MATLAB scripts.

“Forward” (“Source”-to-”Target”) and “backward” (“Target”-to-”Source”) APs propagating along the microchannels were identified based on the extracted spike times. Propagation sequence identifications were performed as previously reported (Costa et al., 2020; Mateus et al., 2021). The custom scripts are available at Github (https://github.com/paulodecastroaguiar/Calculate_APs_velocities_in_MEAs). A propagating event had to fulfill the following requirements: event detected over the entire microchannel (5 electrodes; thus satisfying 4 inter-electrode propagations) and time delay between electrode pairs lower than or equal to 1 ms (minimum propagation velocity of 0.2 m/s).

The method’s validity was confirmed with experiments with tetrodotoxin (TTX) injection (1 µM) on the “Source” compartment for all asymmetric designs (n = 4). TTX injection abolished forward spiking activity (to a fraction of ∼0.001% post-TTX), strongly supporting that the events detected as “forward” are a result of the neuronal activity on the “Source” compartment.

### Network burst detection and propagation characterization

For the detection of NBs, an inter-spike interval (ISI)-statistics-based adaptive threshold alternative, developed by Pasquale et al. (Pasquale et al., 2010), was implemented. After exhaustive analysis, we fine-tuned the parameters to comply with diverse network dynamics. As such, if the percentage of active electrodes was higher or equal to 50%, the values of *minPercElec, maxISITh, maxISImax* were set to 15%, 250 and 600, respectively, and lowered to 10%, 100 and 450, otherwise. *minSpikes* was set to 4, regardless.

After identifying each population’s (“Source” and “Target”) NBs, we assessed the directionality of putative population-to-population communications. If each population initiated concurrent NBs within a [20, 240] ms causality window (as in (Winter-Hjelm et al., 2023)), we considered these events as related. “Forward” events correspond to a “Source” population’s NB driving a “Target” population’s NB and “backward” events to *vice versa*. Each population’s NB could only be considered for a single propagation following sequential order.

### Inclusion criteria and statistical analyses

Electrodes with a mean firing rate (MFR) of at least 0.1 Hz were considered as “active” electrodes. A minimum of 10% of each of the populations’ electrodes (“Source” and “Target”) had to be active for the recording to be considered for downstream analysis. Likewise, to ensure that the populations could communicate, a minimum of 100 summed (forward + backward) propagating events on the microchannels had to be detected. For the comparison of the “forward” and “backward” NB propagation fractions (**Fig. 5D**), a minimum of 3 propagations was considered.

All statistical data is presented as mean ± SD unless otherwise specified. The method of Shapiro-Wilk was used as a normality test and parametric or non-parametric tests were chosen as appropriate. Sample sizes and used tests are indicated in the figures’ legends or the main text. Statistical significance was considered for p < 0.05.

